# Distinct roles of general anesthesia activated CeA neurons in acute versus late phase of neuropathic pain

**DOI:** 10.1101/2024.09.11.612553

**Authors:** Junli Zhao, Kenta Furutani, Aidan McGinnis, Joseph P Mathew, Fan Wang, Ru-Rong Ji

**Author notes:** Correspondence should be addressed to: Ru-Rong Ji Center for Translational Pain Medicine, Department of Anesthesiology, Duke University Medical Center, Durham, North Carolina, NC 27710, USA.

## Abstract

A previous study discovered a distinct population of GABAergic neurons in the <u>ce</u>ntral <u>a</u>mygdala (CeA) that can be activated by <u>g</u>eneral <u>a</u>nesthesia (CeA_GA_) and exert analgesic functions (Hua et al., 2020). To independently reproduce these prior findings and to investigate the electrophysiological properties of CeA_GA_ neurons, we first used 1.2% isoflurane to induce c-Fos activation in the mouse brain and validated the *Fos* expression by RNAscope *in situ* hybridization. Indeed, isoflurane induced robust Fos expression in CeA and these Fos^+^ CeA_GA_ neurons are GABAergic neurons (Vgat^+^). We next used Fos-TRAP2 method (different from the CANE method used in the prior study) to label CeA_GA_ neurons (tdTomato^+^). Our *ex vivo* electrophysiological recordings in brain slices revealed that compared to Fos-negative CeA neurons, CeA_GA_ neurons had significantly higher excitability and exhibited distinct patterns of action potentials. Chemogenetic activation of Fos-TRAPed CeA_GA_ neurons was effective at increasing pain thresholds in naïve mice and mice with early-phase neuropathic pain 2 weeks after spared nerve injury (SNI). However, the same chemogenetic activation of CeA_GA_ neurons only had modest analgesia in the late phase of SNI at 8 weeks, although it was highly effective in reducing chronic pain-associated anxiety behaviors at this stage. We found that Fos-negative CeA neurons, but not CeA_GA_ neurons, exhibited increased excitability in the late-phase of SNI, suggesting that chronic pain causes a shift in the relative activity of the CeA microcircuit. Interestingly, Fos-negative neurons exhibited much higher expression of K^+^-Cl^−^ cotransporter-2 (KCC2), and KCC2 expression was downregulated in the CeA in the late-phase of neuropathic pain. These results support the idea that targeting CeA_GA_ neurons may provide therapeutic benefits for pain relief and chronic pain-associated anxiety. Our findings also suggest distinct roles of CeA_GA_ neurons in regulating physiological pain, acute pain, and chronic pain with a possible involvement of KCC2.

## 1. Introduction

Unlike acute pain that primarily serves a protective role, chronic pain, especially long-lasting chronic pain is a debilitating disease and currently afflicts more than 20% of adults in US. The connections between the widespread burden of chronic pain and the country’s opioid epidemic highlight the urgent need to develop non-addictive pain treatments. Towards this goal, recently a specific subset of <u>ce</u>ntral <u>a</u>mygdala (CeA) neurons activated by <u>g</u>eneral <u>a</u>nesthesia was identified (called CeA_GA_ neurons) to have potent pain-suppression effects^1^. This previous study showed that diverse general anesthetics all induced the expression of the immediate early gene Fos in these CeA_GA_ neurons. Using an activity-dependent targeting method called CANE^2^, it was found that optogenetic activation of these neurons reduced mechanical and thermal sensitivity in sensory tests, stopped ongoing self-caring behaviors in formalin pain models, and abolished mechanical hypersensitivity in a neuropathic pain model (chronic constriction of the infra-orbital nerve)^1^. These findings suggested the intriguing possibility of developing pain-relief therapies by targeting these CeA_GA_ neurons.

To further investigate the potential of CeA_GA_ neurons as targets for pain treatment, we aim to validate the functions using an independent method to specifically target these neurons. Furthermore, the pain-suppression effect of CeA_GA_ neurons in the late phase of long-lasting chronic pain models has not been tested. This issue is important to address from therapeutic perspective. Additionally, the previous study did not characterize the electrophysiological properties of these CeA_GA_ neurons in either naïve mice or in mice with chronic pain conditions. CeA_GA_ neurons are embedded in the complex CeA nucleus with diverse cell types^3–5^. It remains unknown whether CeA_GA_ and other CeA neurons undergo intrinsic property changes in the late phase of persistent pain, which may alter the potency of these neurons in modulating pain.

Here we validate the Fos induction in CeA_GA_ neurons by the anesthetic isoflurane. We successfully employed the Fos-TRAP2 method^6^ (different from the CANE method used in the previous study^1^) to target the CeA_GA_ neurons, and used chemogenetic approaches to test their functions in the spared nerve injury model (SNI) of neuropathic pain. Notably, the SNI model^7^ produces long-lasting pain hypersensitivity, allowing us to examine the effect of CeA_GA_ neurons in both early phase (∼2 weeks) and late phase (>8 weeks) of chronic neuropathic pain. We also performed simultaneous slice physiology studies of labeled CeA_GA_ (tdTomato^+^) and other CeA (unlabeled, tdTomato^-^) neurons in both naïve and SNI mice. Interestingly, we discovered differential effect of CeA_GA_ neurons in early and late phase of SNI, which likely result from electrophysiological property changes.

K^+^-Cl^−^ cotransporter-2 (KCC2) in the spinal cord dorsal horn plays an important in keeping the homeostasis of physiological pain, as KCC2 plays a key role in maintaining the chloride ion gradient, which is necessary for GABAergic synaptic transmission and suppression of pain^8^. In chronic pain conditions such as nerve injury, KCC2 is downregulated in spinal cord neurons, leading to a reversal of the chloride gradient and development of neuropathic pain ^8,9^. We found much higher percentage of Fos-negative neurons expressing KCC2, compared to Fos^+^ neurons. We also observed KCC2 downregulations in the CeA after nerve injury, especially in the late phase of SNI, suggesting a potential role of KCC2 dysregulation in altered CeA population activity in chronic pain.

## 2. Methods

### 2.1 Animals

Adult mice of both sexes (8-26 weeks) were used for behavioral tests and electrophysiological studies. TRAP2 mice (Fos^2A-iCreER^, Stock No: 030323), Ai9 mice (Stock No: 007909) and wild-type (WT) mice (Stock No: 000664) were purchased from Jackson Laboratory and maintained at Duke University Animal facilities. TRAP2; Ai9 mice were generated by crossing TRAP2 mice with Ai9 mice. All transgenic mice were maintained on a C57BL/6 background. All the mouse procedures were approved by the Institutional Animal Care & Use Committee of Duke University. Mice were housed under a 12-hour light/dark cycle with food and water available ad libitum. Animals were randomly assigned to each experimental group. Two to five mice were housed in each cage. All behavioral measurements were conducted during daytime (light cycle). Animal experiments were conducted in accordance with the National Institutes of Health Guide for the Care and Use of Laboratory Animals. The experiments in this study were done and analyzed by researchers who are blinded to genotype or treatment.

### 2.2 Viral vectors

pAAV-hSyn-DIO-hM3D(Gq)-mCherry (cat# 44361-AAV5) and pAAV-hSyn-DIO-mCherry (cat# 50459-AAV5) were produced by Addgene and stored in aliquots at -80°C until use.

### 2.3 Stereotaxic intra-central amygdala injections

For intra-central amygdala injections, mice were anesthetized with isoflurane (4% for induction and 1.5% thereafter for maintenance), and their heads were fixed in a stereotaxic apparatus (David Kopf Instruments). AAV5-hSyn-DIO-mCherry or AAV5-hSyn-DIO-hM3Dq-mcherry was injected into the CeA (–1.24 mm from bregma, ±2.8 mm lateral from midline and 4.5 mm ventral to skull). We injected 300 nL of virus at a rate of 60 nl min^−1^ using the UltraMicroPump injection system (World Precision Instruments). After each injection, the needle was left in place for an additional 10 min for efficient diffusion of the virus-containing reagent and then slowly withdrawn. Behavioral experiments or electrophysiological recordings were performed at least 3 weeks after virus injection. Virus infection was examined at the end of all the tests and only mice that had virus injection restricted to the CeA were used for analysis.

### 2.4 Isoflurane induced Fos expression and labeling

To induce Fos expression in WT mice, mice were put in a camber and anesthetized with isoflurane (1.2% isoflurane and 0.75% oxygen) for 1.5 hour. After the isoflurane induction, mice were transcardially perfused with PBS and 4% paraformaldehyde solution for staining and imaging analysis. To label Fos^+^ neurons in TRAP2 mice;Ai9 mice and TRAP2 mice injected with AAV5-hSyn-DIO-mCherry or AAV5-hSyn-DIO-hM3Dq-mcherry virus, we anesthetized mice with isoflurane for 1 hour, then injected mice with 50 mg/kg 4-Hydroxytamoxofen and further anesthetized mice with isoflurane for another 3 hours. The 4-Hydroxytamoxofen (Sigma, Cat# H6278) was dissolved in 100% ethanol as 20 mg/mL stock solution and kept in –80°C. On the experiment day, we added corn oil to further dissolve 4-Hydroxytamoxofen to 10 mg/mL and sonicate until solution cloudiness clears. Then the solution was vacuum centrifuged for 10 min to evaporate the alcohol from the final injection solution. In TRAP2;Ai9, robust tdTomato expression was seen 10 days after isoflurane induction. In the TRAP2 mice injected with AAV5-hSyn-DIO-mCherry or AAV5-hSyn-DIO-hM3Dq-mcherry virus, robust labeling was seen 4 weeks after isoflurane induction.

### 2.5 Chemogenetic manipulation

For chemogenetic manipulation of the Fos^+^ neurons in CeA, TRAP2 mice were injected with AAV5-hSyn-DIO-mCherry or AAV5-hSyn-DIO-hM3Dq-mcherry. Three weeks later, these mice were injected with 4-Hydroxytamoxofen and anesthetized with isoflurane to express Cre-dependent DREADDs. CNO (3 mg/kg, i.p, Item No. 25780, Cayman Chemical) was injected 30 min before doing behavioral experiments and data were collected between 1-6 hours post-injection.

### 2.6 Spared nerve injury (SNI) model of neuropathic pain

The SNI model was induced as previously reported^7,10,11^. Mice were anesthetized with isoflurane (4% for induction and 1.5% thereafter for maintenance), and the skin of the lateral left thigh was incised. The cranial and caudal parts of the biceps femoris muscle were separated and held apart by a retractor to expose the sciatic nerve and its three terminal branches: the sural, common peroneal and tibial nerves. The procedure involved tightly binding and ligation of the tibial and common peroneal nerves with 6/0 silk sutures, followed by the removal of a small segment (1-2 mm) of the nerve just distal to the ligation site. Any stretching or contact with the intact sural nerve was avoided. Following the nerve injury, the muscle layer was carefully closed, and the skin was stitched using hidden sutures to prevent wound reopening from biting. For the sham control group, a similar exposure of the sciatic nerve and its branches was performed, but without any ligation or transection.

### 2.7 Nociceptive testing

#### Von Frey Filaments Test

In Von Frey test, mice underwent a habituation process for three days before baseline measurements. Mice were placed in individual plastic chambers on a wire mesh during both the habituation and testing phases, allowing their plantar sites to be easily accessed. The evaluation of mechanical allodynia involved using Von Frey filaments, with weights varying from 0.008 to 2.0 g (Stoelting Co.), applied to the hindpaw’s plantar surface. Each filament was pressed down just enough to cause a slight bend and was held in place for around 2 seconds to elicit a response. If the mouse did not respond within 2 seconds, the next test involved a heavier filament. In contrast, a reaction led to testing with a lighter filament. This procedure was repeated to gather six measurements per mouse or until a pattern of four consecutive positive or negative responses emerged. The up-down method was used to calculate the 50% withdrawal threshold^12^. For the assessment of mechanical hyperalgesia, the frequency of responses was determined. This was done by applying a 0.4 g filament to the plantar site ten times. The response frequency was then calculated with the formula: (total number of withdrawals / 10) × 100%.

#### Hargreaves Test

Assessment of thermal sensitivity in mice was performed using the Hargreaves test^13^. Mice were tested in ventilated opaque white Plexiglas testing chambers placed on an elevated platform with a clear glass surface. Following a 1-hour period of habituation, a thermal stimulus from a constant radiant heat source was delivered through the glass bottom of the chamber to the plantar surface of the hindpaw (IITC Life Sciences). The time to paw withdrawal was recorded. The average of three latencies were taken from each hindpaw.

#### Acetone Evaporative Test

The acetone evaporative test was adapted to measure sensitivity to a cold stimulus. On both habituation and testing days, the mice were placed in plastic chambers set atop a wire mesh table, allowing access to their plantar sites. Acetone (Sigma) was drawn into a 20 µL syringe and a drop was lightly applied through the wire mesh to the plantar surface of the hindpaw, being careful not to directly touch the paw with the syringe to avoid false withdrawal responses. Following acetone application, nociceptive responses were video-recorded and the total time of lifting, licking, and/or were quantified.

### 2.8 Elevated plus maze test

To evaluate the impact of chemogenetic manipulation of CeA_GA_ neurons on anxiety-like behaviors, we conducted the elevated plus maze. The apparatus (Stoelting Co.) is comprised of two open arms (35 × 5 cm^2^) and two closed arms (35 × 5 × 15 cm^3^) elevated 50 cm above the ground. On the day of the experiment, each mouse was initially explore the environment. Mouse behaviors, including distance in the open arms, time spent in the open arms and entries in the open arms were tracked and recorded using ANY-maze software (Stoelting Co.).

### 2.9 Open filed test

To assess anxiety-like behavior resulting from chemogenetic manipulation of the CeA_GA_ neurons, we employed the open field test. This test was conducted in a square arena measuring 45 × 45 × 45 cm^3^ (TAP plastics) and lasted for 10 minutes. We defined the center of the arena as a square area encompassing 50% of the total area. Metrics such as the distance traveled in the center square, time spent in center square, and the number of entries into center square were automatically recorded and analyzed by the ANY-maze software (Stoelting Co.).

### 2.10 Novel object recognition test

To assess cognitive behavior following chemogenetic manipulation of the CeA_GA_ neurons, we utilized the novel object recognition test^14^. On the first day of the experiment, mice were placed in a 30 × 30 × 30 cm^3^ square arena (TAP plastics) for habituation. On the following day, two identical objects were positioned in separate corners of the arena. After this, the mice were returned to their home cages for a 30 min retention interval. Subsequently, one of the identical objects was replaced with a new one for the novel object recognition test. Post-retention interval, the animals were reintroduced into the arena for an additional 5 min of exploration. Valid exploration was defined as the mouse either touching an object with its nose or paying attention to it from a distance of less than 1 cm. Actions such as turning around, climbing, or sitting on the object were deemed invalid. Exploration time for the familiar or novel object was evaluated using a discrimination index, calculated as DI = (T_N_ − T_F_)/(T_N_ + T_F_) × 100%. All animal behaviors were video-recorded and analyzed by an experimenter blind to the testing conditions.

### 2.11 Tail suspension test

To evaluate depression-like behavior in response to chemogenetic manipulation of the CeA_GA_ neurons, we employed the tail suspension test. In this test, mice were suspended by their tails using adhesive tape, placed approximately 1 cm from the tip of the tail and ∼15 cm above the table surface. To prevent the animals from climbing or hanging onto their tails, plastic tubes were fitted over the tails. The mice were then video recorded for a duration of 6 minutes. A mouse was deemed immobile only if it remained passive and motionless for a minimum of 2 seconds. An experimenter, blinded to the testing conditions, quantified the time each mouse spent

### 2.12 Forced swimming test

To evaluate depression-like behavior in response to chemogenetic manipulation of the CeA_GA_ neurons, the forced swimming test was also utilized. In this test, mice were placed in a glass container filled with water to a depth of 20 cm, maintained at a temperature range of 22-24°C. The test lasted for 6 minutes, during which the mice were video recorded. After the test, the wet mice were gently dried with a towel and then placed in a drying cage with a heat lamp positioned above and a heat pad underneath. This ensured that the mice were comfortably warmed and dried. A mouse was considered “immobile” if it floated with minimal movement, only enough to keep its nose above water. An experimenter, who was blind to the testing conditions, quantified the time each mouse spent immobile.

### 2.13 Mouse brain slice preparation and patch-clamp recording

Mice were anesthetized with isoflurane and subsequently decapitated. Their brains were quickly transferred to an ice-cold, artificial, sucrose-based cerebrospinal fluid (ACSF) solution. This solution contained the following (in mM): 75 Sucrose, 87 NaCl, 2.5 KCl, 1.25 NaH_2_PO_4_, 0.5 CaCl_2_, 7 MgCl_2_, 26 NaHCO_3_, and 25 glucose. Brain slices that included the CeA region, each 350 μm thick, were prepared using a Leica VT1200S vibratome. These slices were then transferred to a normal ACSF solution, composed of (in mM): 124 NaCl, 3 KCl, 1.25 NaH_2_PO_4_, 2 CaCl_2_, 1 MgSO_4_, 26 NaHCO_3_, and 10 glucose, and maintained at 34°C. All extracellular solutions were constantly carbogenated with a mixture of 95% O_2_ and 5% CO_2_. Generally, the slices were kept in ACSF for at least 1 hour before any recordings were made.

Whole-cell recordings were conducted at a controlled temperature of 34 ± 1°C, using an automatic temperature controller from Warner Instruments and an EPC10 amplifier from HEKA. The CeA was identified by its distinctive fiber bundles, which clearly delineate its structure. Both tdTomato^+^ and tdTomato^-^ neurons were recorded and confirmed through tdTomato fluorescence in TRAP2; Ai9 mice. Data were low-pass filtered at 2 kHz and sampled at 10 kHz. The patch pipettes used were filled with a solution containing the following (in mM): 135 K-gluconate, 5 KCl, 0.5 CaCl_2_, 2 MgCl_2_, 5 HEPE, 5 EGTA, and 5 MgATP, adjusted to a pH of 7.3 and an osmolarity of 290–300 mOsm/L. When filled with this solution, the pipettes had a resistance of 4-8 MΩ. Resting membrane potentials (RMPs), spontaneous spikes, rheobase, and action potentials were recorded in current-clamp mode. Rheobase and action potentials were evoked by current injection steps ranging from 0 to 130 pA, with 10 pA increments. Data analysis was performed using Patchmaster software from HEKA.

### 2.14 *In situ* hybridization in mouse brain sections

Mice were deeply anesthetized using isoflurane and then perfused first with PBS, followed by 4% paraformaldehyde (PFA). After perfusion, the brains were extracted and post-fixed overnight at 4°C in 4% PFA. Following post-fixation, the brains underwent dehydration in 30% sucrose solution. Subsequently, all tissues were embedded in OCT medium. The embedded tissues were then cryosectioned to 30-μm thick tissue sections, which were mounted onto charged slides. *In situ* hybridization was performed using the RNAscope® Multiplex Fluorescent Reagent Kit v2 from Advanced Cell Diagnostics (Cat# 323100), following the manufacturer’s instructions. Probes specific to mouse genes *Fos* (Cat# 316921), *Slc17a6* (Cat# 428871), *Slc32a1* (Cat# 319198), *Prkcd* (Cat# 441791), and *Sst* (Cat# 404631) were utilized.

### 2.15 Immunochemistry in mouse brain sections

The brain samples were prepared as shown in Session 2.15. Free-floating brain sections (30 μm) were cut in a cryostat (Leica CM1950). Tissue sections were washed several times in PBS and blocked with 0.1% Triton X-100 and 10% goat serum for 1 h at room temperature. The sections were then incubated overnight at 4°C in a humidified chamber with the following primary antibodies: anti-KCC2 antibody (rabbit, 1:100, Sigma, Cat# 07-432). After washing, the sections were incubated with Nissl/Neuro Tracer 640/660 (1:100, ThermoFisher Scientific, Cat# N21483) or species-specific secondary antibodies conjugated to 488-nm (1:500, ThermoFisher Scientific) for 2 h at room temperature. Sections were subsequently washed and coverslipped using Fluoroshield™ with DAPI (Sigma, Cat# F6057).

### 2.16 Image acquisition and analysis

The stained sections were examined using a Zeiss LSM 880 confocal microscope, employing both Z-stack and Tile Scan features. Image analysis was exclusively conducted on images captured with the 20X objective. We utilized sequential acquisition across multiple channels, with z-stacks collected in 1.0 μm steps. The image stacks were then processed into maximum intensity z-projections and stitched together using Zeiss Zen software. Quantitative analysis was focused on the CeA, specifically between bregmas -1.06 and -1.46. In each mouse, three sections were analyzed, and three to four mice per group were included in the analysis. To ensure consistency, laser intensity, gain, and pinhole settings were maintained constant across experiments. All analyses and quantifications were performed by researchers blinded to the experimental conditions.

All data are presented as the mean ± SEM. Details regarding the sample size and statistical analysis for each experiment are provided in the figures and figure legends. In electrophysiology studies, each data point represents an individual neuron. Neurons were sampled from a minimum of three different animals for all analyses. In behavioral studies, each data point corresponds to an individual animal. Unless specified otherwise in the figure legends, behavioral, biochemical, and some electrophysiological data were analyzed using either two-way or one-way ANOVA, accompanied by Bonferroni’s or Tukey’s post-hoc test for multiple comparisons. Differences between two groups were compared using a two-tailed t-test. The threshold for statistical significance was set at *P* < 0.05. Asterisks in the figures denote significance levels as follows: \**P* < 0.05, \*\**P* < 0.01, \*\*\**P* < 0.001, \*\*\*\**P* < 0.0001. Statistical analyses were conducted using Prism GraphPad version 10.0.

## 3. Results

### 3.1 Isoflurane induces Fos activation in subsets of GABAergic neurons in CeA and glutamatergic neurons in PVT, PVN and SON

We started by independently validating the presence of CeA_GA_ neurons^1^. We induced general anesthesia (GA) using 1.2% isoflurane for 1.5 hours and collected mouse brains. We examined the *Fos* expression by RNAscope *in situ* hybridization (ISH) using a highly sensitive and selective commercial probe (Fig. 1A). Our results showed that the central amygdala (CeA) is one of the most strongly activated brain region by isoflurane (Fig. 1B, C). Additionally, we found strong *Fos* expression in the paraventricular nucleus of the thalamus (PVT), the paraventricular nucleus of the hypothalamus (PVN), and the supraoptic nucleus (SON) in the hypothalamus^15^ (Fig. 1B, Fig. S1A-S1B). In our negative control experiment, in which animals were only exposed to 0.75% oxygen for 1.5 hours (Fig 1A), we found limited *Fos* expression in the CeA (Figs. 2A, B, Fig. S1C). Furthermore, more Fos^+^ neurons were found in the posterior region of CeA compared to oxygen control (Fig. 2B).

**Figure 1.**
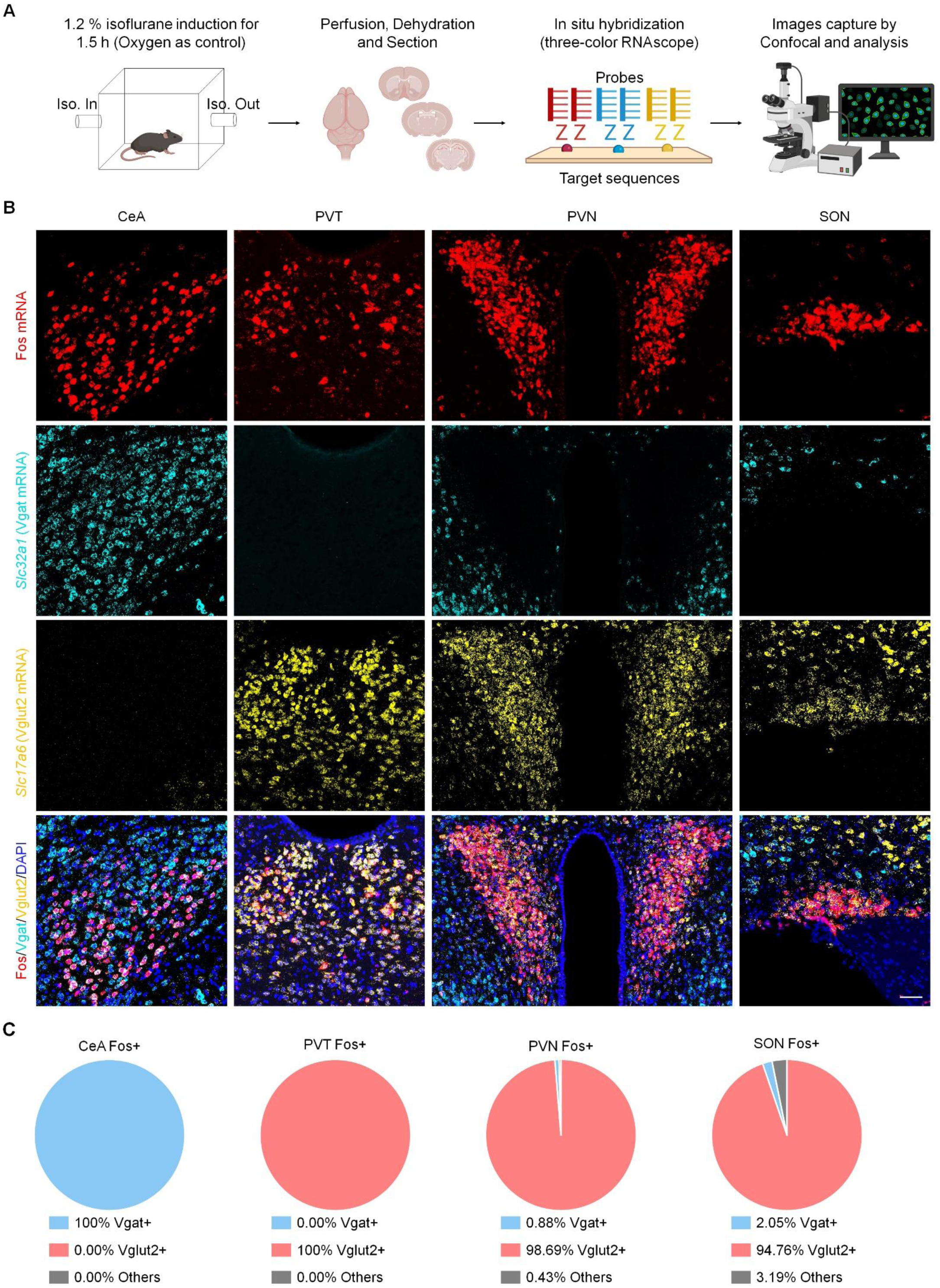
Isoflurane induces Fos activation mainly in GABAergic neurons in CeA and glutamatergic neurons in PVT, PVN and SON. **(A)** Schematic of experimental design for *in situ* hybridization in mouse brain slices. Scale bar: 50 µm. **(B)** Representative image of *Fos* (red), *Vgat* (*Slc32a1*, cyan) and *Vglut2* (*Slc17a6*, yellow) expression in CeA, PVT, PVN and SON regions. **(C)** Quantification of the percentage of *Fos^+^* neurons over *Vgat^+^*, *Vglut2^+^* or other neurons in CeA, PVT, PVN and SON regions.

**Figure 2.**
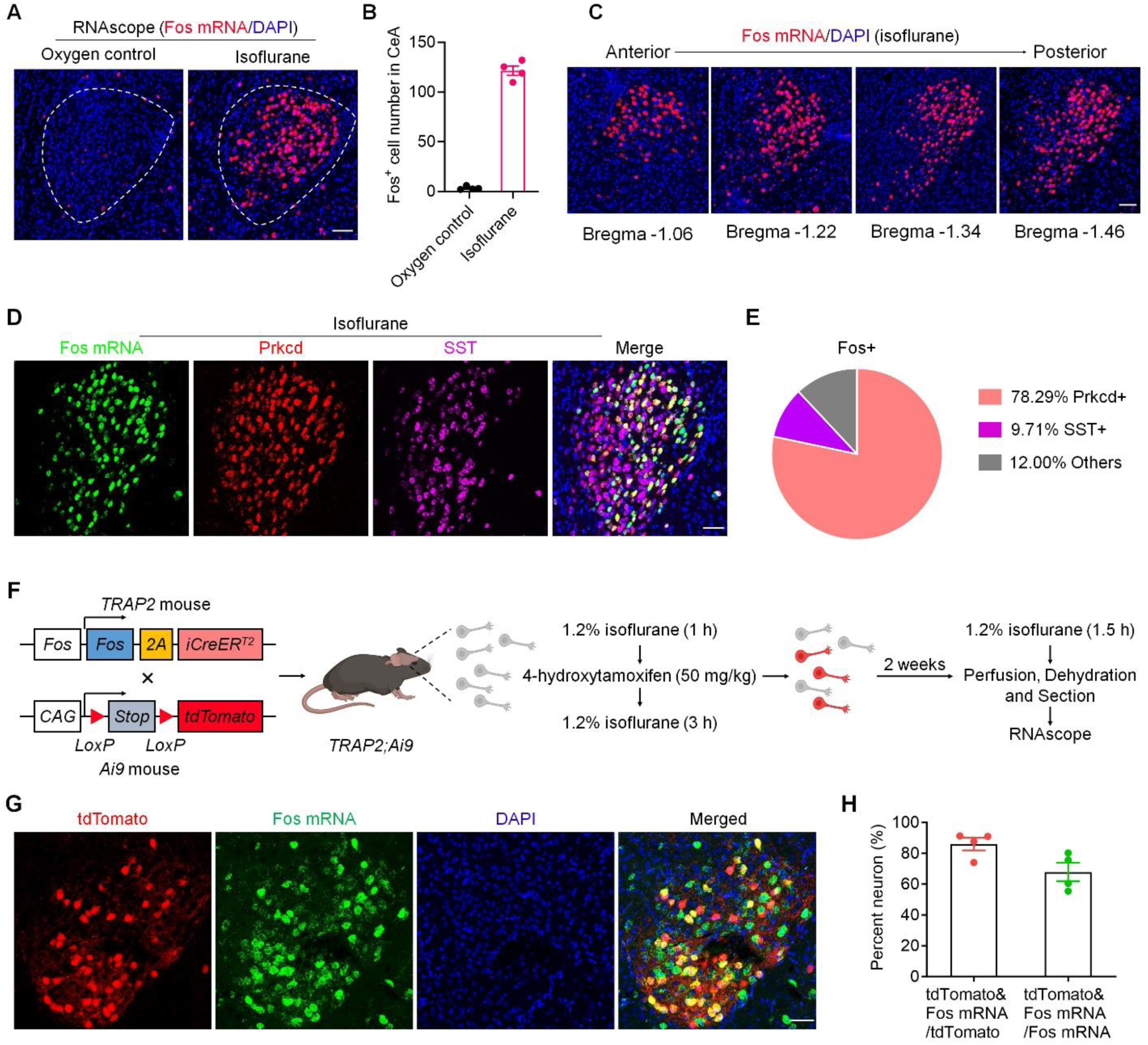
Characterization of isoflurane induced Fos activation in the CeA region of WT mice and TRAP2; Ai9 mice. **(A)** Representative images of *Fos* mRNA expression in WT mice after oxygen control and isoflurane exposure. Scale bar: 50 µm. **(B)** Quantification of averaged *Fos^+^*cells in the CeA after oxygen (control) and isoflurane exposure. **(C)** Representative images showing *Fos* expression induced by isoflurane from Bregma −1.06 mm to −1.46 mm of WT mice. Scale bar: 50 µm. **(D)** Representative images showing the expression of *Fos*, *PKCδ* (*prkcd*) and *SST* in WT CeA regions after isoflurane induction. Scale bar: 50 µm. **(E)** Percentage of *PKCδ^+^* and *SST^+^* neurons over *Fos^+^* neurons in WT CeA regions after isoflurane exposure. **(F)** Schematic of experimental design for trapping *Fos^+^* CeA_GA_ neurons using TRAP2; Ai9 mice. **(G)** Representative images of tdTomato^+^ and *Fos^+^* neurons in TRAP2; Ai9 mice after isoflurane exposure. Scale bar: 50 µm. **(H)** Quantification of the percentage of tdTomato^+^/*Fos^+^* neurons over tdTomato^+^, or *Fos^+^* neurons in TRAP2; Ai9 mice CeA region after isoflurane exposure. All data are presented as the mean ± SEM.

To further examine whether the GA-activated Fos^+^ neurons are excitatory or inhibitory in these brain regions, we conducted triple staining of ISH using RNAscope probes for *Fos* mRNA, *Vgat* mRNA (*Slc32a1* to label inhibitory neurons), and *Vglut2* mRNA (*Slc17a6* to label excitatory neurons) in the CeA, PVT, PVN and SON. We further quantified the percentage of *Fos^+^* neurons in the *Vgat^+^*and *Vglut2^+^* populations in the brain regions. Consistent with known literature and the recent publication^1^, *Fos^+^* CeA_GA_ neurons are 100% inhibitory (Fig. 1B, C), whereas *Fos^+^* PVT neurons are 100% excitatory (Fig. 1C), and 98.7% PVN neurons and 94.8% SON neurons expressed *Vglut2* (Fig. 1C). Furthermore, 78.3% and 9.7% of Fos^+^ neurons in the CeA express *PKCδ* (*prkcd*) and *somatostatin* (*Sst*), respectively (Fig. 2D, E). These results validated the previous findings and promoted us to use a Fos-based method to target CeA_GA_ neurons for further investigations.

### 3.2 Electrophysiological characterization of CeA_GA_ neurons using Fos-TRAP2 based targeting in naïve mice

The previous study used CANE method^1^ to label and manipulate CeA_GA_ neurons. Here we first tested whether these neurons can be efficiently and specifically labeled using a different activity-dependent method, Fos-TRAP2^6^. We crossed Fos-TRAP2 mice with Ai9 tomato-reporter mice^16^ (Fig. 2F). To trap CeA_GA_ neurons, we subjected these mice to 1.2% isoflurane for 1 hour, at which point we injected 4-hydroxytamoxifen (4-OHT, 50 mg/kg) into the mice, followed by additional 3 hours of isoflurane anesthesia. 2 weeks later, we re-exposed mice to isoflurane to re-induce *Fos*. ISH analysis revealed a high ratio overlap between *Fos* mRNA^+^ and tdTomato^+^ neurons in the CeA region of Fos-TRPA2; Ai9 (Fig. 2G). About 85% of tdTomato^+^ neurons express *Fos* and about 65% *Fos^+^* neurons are tdTomato^+^ (Fig. 2H). Thus Fos-TRAP2 method can selectively label CeA_GA_ neurons with good efficiency. We therefore use this method to investigate the electrophysiological properties of these neurons and compared results to other unlabeled CeA neurons.

Two weeks after trapping CeA_GA_ neurons in Fos-TRPA2; Ai9 mice, we prepared acute brain slices for patch-clamp recordings (Fig. 3A). Under the bright-field microscope, the cell bodies of CeA neurons in brain slices could be clearly visualized (Fig. 3B). Furthermore, tdTomato^+^ (CeA_GA_) and negative neurons (non-CeA_GA_) were identified for patch-clamp recordings (Fig. 3C). We found differences in firing patterns of tdTomato^+^ and tdTomato^-^ negative neurons in CeA (Fig. 3D, E). Characterization of 34 tdTomato^+^ CeA_GA_ neurons revealed two firing patterns: regular spiking (RS) in 23 neurons (68%) and late-spiking (LS) in 11 neurons (32%). Recordings in 34 tdTomato^-^ non-CeA_GA_ neurons revealed three firing patterns: regular spiking (RS) in 15 neurons (44%), late-spiking (LS) in 15 neurons (44%), and low threshold bursting (LTB) in 4 neurons (12%). Thus, CeA_GA_ consists of higher proportions of RS neurons, compared to other CeA neurons (68% vs. 32%), but no LTB type of neurons. Many CeA_GA_ neurons express *Prkcd* and a previous study found that *Prkcd^+^* CeA neurons are of RS and LS types^3,5^, consistent with our results. Furthermore, we found that CeA_GA_ neurons exhibited less negative resting membrane potential (RMP, Fig. 3F), lower rheobase (Fig. 3G), and higher numbers of action potentials following current injection (Fig. 3H, I). These data suggest that Fos-TRAPed CeA_GA_ neurons have higher excitability than other CeA neurons in naïve mice. Additional examination in RS (Fig. 3J, K) and LS neurons (Fig. 3L, M) also showed greater excitability of CeA_GA_ neurons.

**Figure 3.**
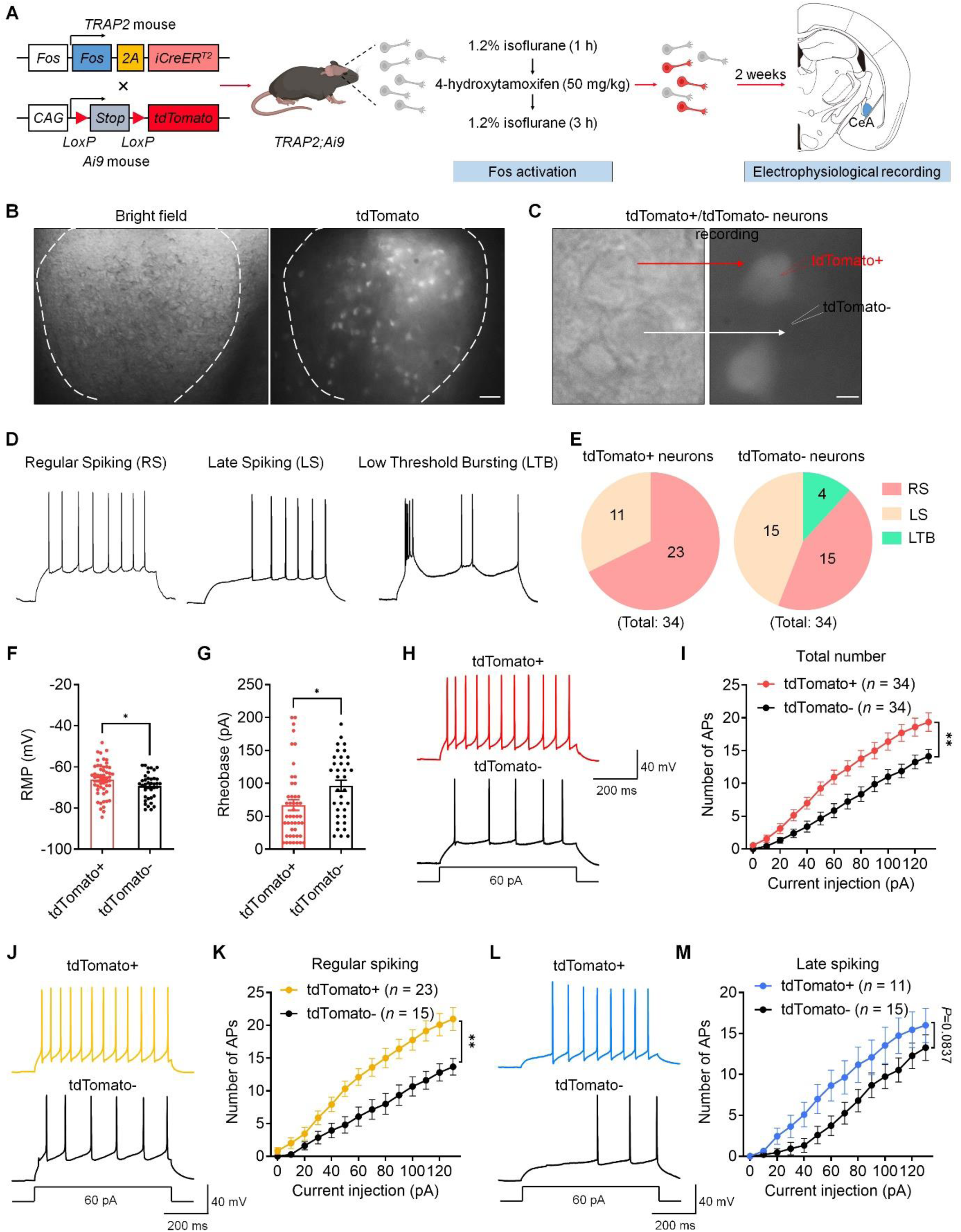
CeA_GA_ neurons have higher excitability than non-CeA_GA_ neurons in the CeA region in naïve mice. **(A)** Schematic of experimental design for labeling CeA_GA_ neurons with tdTomato and electrophysiological recordings in CeA slices of TRAP2; Ai9 mice. **(B)** Representative images showing the CeA region by distinct fiber bundles outlining the nuclei revealed by differential interference contrast microscopy (left) and tdTomato^+^ cells revealed by fluorescent microscopy (right). Scale bar: 50 µm. **(C)** Representative images of tdTomato^+^ and tdTomato^-^ neurons for electrophysiological recording. Scale bar: 10 µm. **(D)** Representative voltage recordings of neurons with regular spiking (RS), late spiking (LS), and low threshold bursting (LTB) in tdTomato^+^ and tdTomato^-^ neurons. **(E)** Proportion of each firing pattern within tdTomato^+^ and tdTomato^-^ populations. **(F)** RMP recorded from tdTomato^+^ and tdTomato^-^ neurons. **(G)** Rheobase is evoked by step current injection and recorded from tdTomato^+^ and tdTomato^-^ neurons. **(H)** Representative AP traces evoked by 60 pA current in tdTomato^+^ and tdTomato^-^ neurons. **(I)** Number of APs evoked by step current injection in tdTomato^+^ and tdTomato^-^ neurons. **(J)** Representative AP traces evoked by 60 pA current in tdTomato^+^ and tdTomato^-^ neurons with regular spiking. **(K)** Number of APs evoked by step current injection in tdTomato^+^ and tdTomato^-^ neurons with regular spiking. **(L)** Representative AP traces evoked by 60 pA current in tdTomato^+^ and tdTomato^-^ neurons with late spiking. **(M)** Number of APs evoked by step current injection tdTomato^+^ and tdTomato^-^ neurons with late spiking. All data are presented as the mean ± SEM. \**P* < 0.05, \*\**P* < 0.01. Unpaired t-test (F, G) or two-way ANOVA (I, K, M).

### 3.3 Chemogenetic activation of CeA_GA_ neurons inhibits mechanical pain in naïve mice and mice with early-phase neuropathic pain

The previous study employed optogenetic stimulation of CANE-captured CeA_GA_ neurons^1^, and here we want to examine whether chemogenetic activation of CeA_GA_ neurons can also modulate pain. We used the Fos-TRAP2 method together with Cre-dependent AAV to express the excitatory DREADDs (hM3Dq) selectively in CeA_GA_ neurons (Fig. 4A-4B). We validated hM3Dq-mCherry expression in CeA_GA_ (Fig. 4C). We tested mechanical and thermal sensitivity in naïve mice before (baseline) and 1 h after intraperitoneal CNO injection (Fig. 4D-G). Von Frey testing showed that CNO treatment significantly increased paw withdrawal threshold (PWT, *P*<0.001, Fig. 4D) and decreased paw withdrawal frequency (PWF, *P*<0.01) to a 0.4 g filament (Fig. 4E). In control mice expressing mCherry AAV, the same CNO treatment had no significant effects on PWT and PWF (*P*>0.05, Fig. 4D, E). Hargreaves radiant heat test showed a mild but significant increase in withdrawal latency (*P* <0.05, Fig. 4F) following CNO treatment in hM3Dq but not control mice, suggesting a moderate inhibition of heat pain.

**Figure 4.**
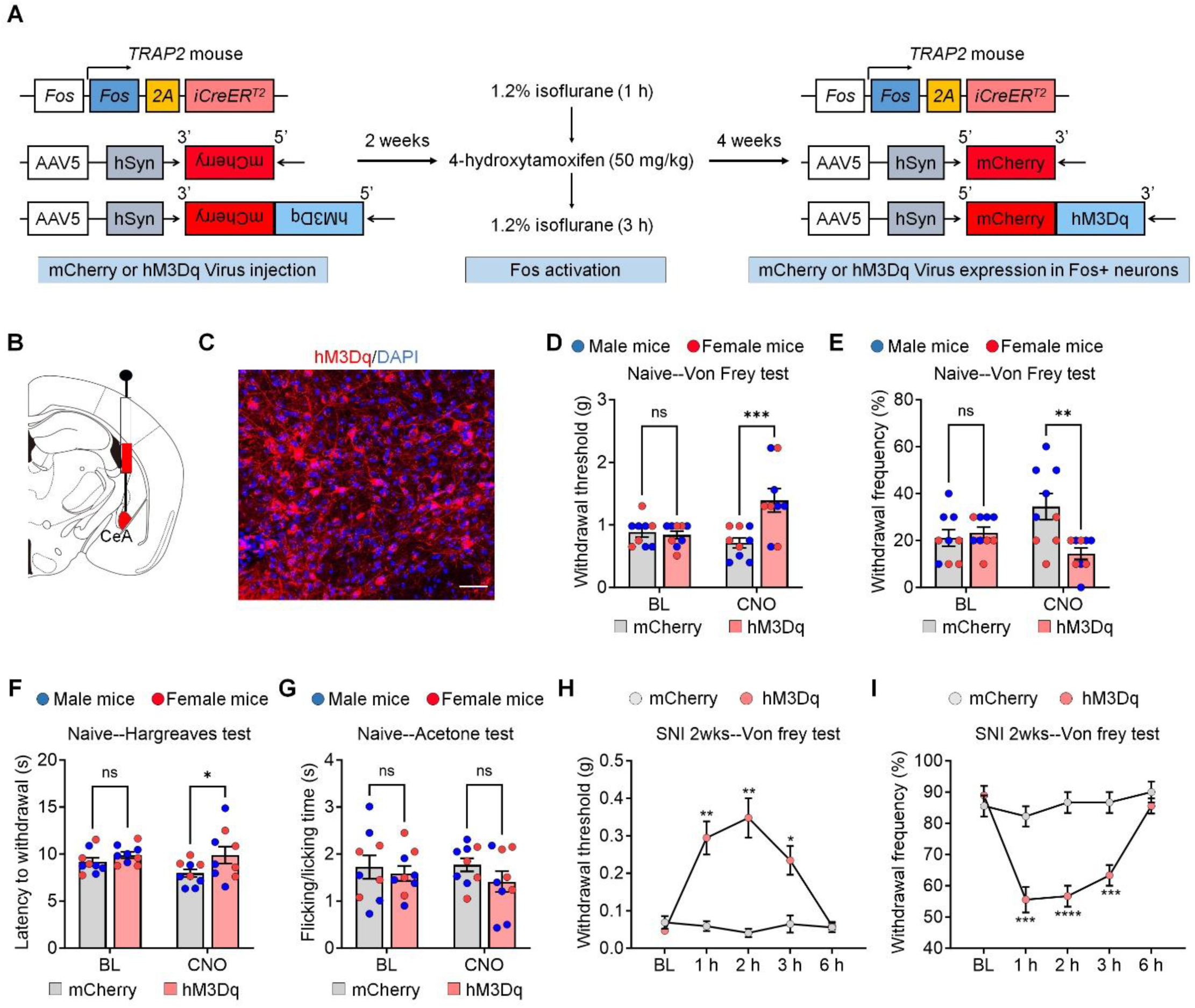
Chemogenetic activation of isoflurane-TRAPed CeA_GA_ neurons attenuates pain in naïve mice and mice with early-phase SNI. **(A)** Illustration of strategy to express excitatory DREADDs (hM3Dq) selectively in isoflurane-TRAPed CeA_GA_ neurons and experimental timeline. **(B)** Schematic of AAV administration in the CeA region of TRAP2 mice. **(C)** Representative image of AAV injection in the CeA region of TRAP2 mice. Red: hM3Dq virus, Blue: DAPI. **(D, E)** Von Frey test showing withdrawal threshold (D) and frequency (E) in naïve mice before (baseline) and 1 h after i.p. CNO injection. **(F)** Hargreaves test showing withdrawal latency in naïve mice before (baseline) and 1 h after CNO i.p. injection. **(G)** Acetone evaporative test showing the total flicking/licking time in naïve mice before (baseline) and 1 h after CNO i.p. injection. **(H)** Von Frey test showing withdrawal threshold in the hindpaw ipsilateral to nerve injury mice before (baseline), and 1 h, 2 h, 3 h, 6 h after CNO i.p. injection. **(I)** Von Frey test showing withdrawal frequency in the hindpaw ipsilateral to nerve injury All data are presented as the mean ± SEM. \**P* < 0.05, \*\**P* < 0.01, \*\*\**P* < 0.001, \*\*\*\**P* < 0.0001. ns: not significant. Two-way ANOVA, accompanied by Bonferroni’s post-hoc test for multiple comparisons (D, E, F, G, H, I).

Acetone test showed no significant effect in cold pain (Fig. 4G). Overall, chemogenetic activation of CeA_GA_ neurons showed analgesic effect in naïve mice, although the effect appear less potent than previously observed with optogenetic activation^1^, likely because hM3Dq-DREADD only moderately activates neurons.

Next, we examined the impact of CeA_GA_ neuron chemogenetic activation in a neuropathic pain model, the spared nerve injury (SNI) model^7^, which was not tested before. Two weeks after SNI surgery, we observed robust mechanical allodynia, as indicated by substantial reduction in PWT (Fig. 4H) and increase in PWF (Fig. 4I), as compared to naïve mice (Fig. 4D, E). A time course study showed that CNO caused significant increase in PWT at 1 h, 2 h, and 3 h but not 6 h, as CNO is likely to be metabolized after 6 h (Fig. 4D). Similarly, CNO injection resulted in significant decrease in PWF at 1 h, 2 h, and 3 h but not 6 h (Fig. 4E). Collectively, these results indicate that chemogenetic activation of CeA_GA_ neurons is highly effective in reducing early phase neuropathic pain.

### 3.4 Chemogenetic activation of CeA_GA_ neurons differentially regulates anxiety, neuropathic pain, and depression-like behaviors in late-phase of nerve injury

Long-term chronic pain is associated with multiple co-morbidities, including anxiety, depression, and cognitive impairment^17–19^. We examined the effect of chemogenetic activation of CeA_GA_ neurons in the late-phase of neuropathic pain, at 8 weeks after SNI surgery (Fig. 5A). CNO (3 mg/kg, i.p.), same concentration as used for early-phase SNI experiments, was injected 30 min before behavioral tests. Fig. 5B shows a representative occupancy heatmap of spatial location in the elevated plus maze (EPM) test for anxiety-like behavior. Compared to control mCherry-mice, chemogenetic activation of hM3Dq-CeA_GA_ mice resulted in significant increases (*P*<0.01) in travel distance in open arms (Fig. 5C), time spent in open arms (Fig. 5D), and entries in open arms (Fig. 5E).

**Figure 5.**
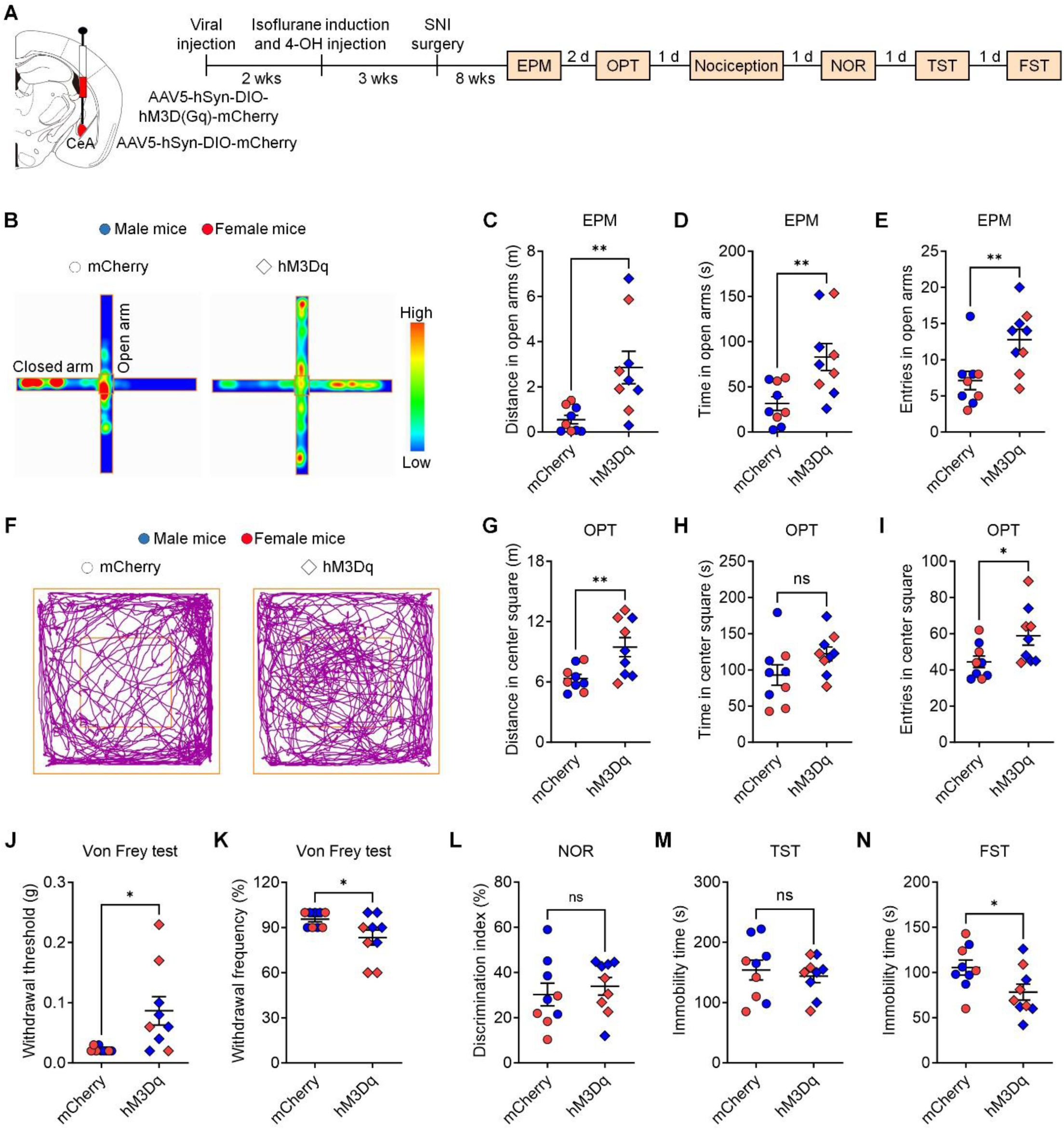
Chemogenetic activation of isoflurane-TRAPed CeA_GA_ neurons decreases chronic pain related anxiety-like behavior. **(A)** Illustration of strategy to express excitatory DREADDs (hM3Dq) selectively in isoflurane-TRAPed CeA_GA_ neurons and experimental timeline in nerve injury model. CNO (3 mg/kg, i.p.) was injected 30 min before the testing. **(B)** Representative occupancy heatmap showing spatial location in the elevated plus maze of a SNI control mouse (mCherry) and a SNI mouse with chemogenetic activation of CeA_GA_ neurons (hM3Dq). **(C-E)** Anxiety-like behavior tested by elevated plus maze for distance in open arms (C), time in open arms (D) and entries in open arms (E) from SNI control mice (mCherry) and SNI mice with chemogenetic activation of CeA_GA_ neurons (hM3Dq). **(F)** Representative travelled traces showing spatial location in the open filed test of a SNI control mouse (mCherry) and a SNI mouse with chemogenetic activation of CeA_GA_ neurons (hM3Dq). **(G-I)** Anxiety-like behavior tested by open field for distance in center square (G), time in center square (H) and entries in center square (I) from SNI control mice (mCherry) and SNI mice with chemogenetic activation of CeA_GA_ neurons (hM3Dq). **(J)** Von Frey test showing withdrawal threshold in SNI control mice (mCherry) and SNI mice with chemogenetic activation of CeA_GA_ neurons (hM3Dq). **(K)** Von Frey test showing withdrawal frequency in SNI control mice (mCherry) and SNI mice with chemogenetic activation of CeA_GA_ neurons (hM3Dq). **(L)** NOR test showing discrimination index of SNI control mice (mCherry) and SNI mice with chemogenetic activation of CeA_GA_ neurons (hM3Dq). **(M)** Depressive-like behavior measured by tail suspension for immobility time in SNI control mice (mCherry) and SNI mice with chemogenetic activation of CeA_GA_ neurons (hM3Dq). **(N)** Depressive-like behavior measured by forced swimming for immobility time in SNI control mice (mCherry) and SNI mice with chemogenetic activation of CeA_GA_ neurons (hM3Dq). All data are presented as the mean ± SEM. \**P* < 0.05, \*\**P* < 0.01. ns: not significant. Unpaired two-tailed t-test (G, H, I, J, K, L, M, N).

We also investigated additional anxiety-like behavior in SNI animals in open filed testing (OPT, Fig. 5F-I). Fig. 5F shows representative traces and spatial location of animals in the open filed test. Notably, chemogenetic activation caused significant increases in distance traveled in center square (*P*<0.01, Fig. 5G) and entries in center square (*P*<0.05, Fig. 5I), but not in time in center square (Fig. 5H).

Von Frey test showed significance changes in PWT (*P*<0.05, Fig. 5J) and PWF (*P*<0.05, Fig. 5K) following the chemogenetic activation of CeA_GA_ neurons in mice with 8-weeks of SNI. However, these changes in mechanical pain were modest, and there was only 10% reduction in PWF (from 90% to 80% response, Fig. 5K), suggesting a reduced analgesic efficacy of chemogenetic activation of CeA_GA_ neurons in late-phase neuropathic pain.

Novel object recognition (NOR) test revealed an impairment in animal’s learning ability 8 weeks after SNI, showing a discrimination index of 30% (Fig. 5L), which is much lower than that of normal naïve mice (∼50%)^14^. CNO treatment did not improve NOR index in the late-phase (Fig. 5L). Thus, the CeA microcircuit may not be involved in regulating cognitive impairment in chronic pain.

Additionally, we examined chronic pain associated depression-like behavior using tail suspension test (TST) and forced swimming test (FST) in mice after 8-week SNI (Fig. 5M, N). We found that CeA_GA_ neuron activation by CNO significantly decreased the immobility time in forced swimming test (*P*<0.05, Fig. 5N) but not in tail suspension test (Fig. 5M). Collectively, these findings suggest that in chronic phase of nerve injury activation of CeA_GA_ neurons has profound effects on anxiety and mild effects on neuropathic pain and depression.

### 3.5 Nerve injury causes significant changes in electrophysiological properties in CeA neurons

To explore the potential cellular mechanisms underlying the decreased efficacy of pain modulation by CeA_GA_ neurons in the late-phase SNI, we compared firing patterns and excitability of tdTomato^+^ (CeA_GA_) and tdTomato^-^ (non-CeA_GA_) neurons in brain slices obtained from Fos-TRPA2; Ai9 mice with SNI-8w (Fig. 6A). In addition to regular spiking (RS) neurons, late spiking (LS), and low threshold bursting (LTB) found in the normal conditions (Fig. 3E), we observed the emergence of another population of neurons with spontaneous spiking (SS, Fig. 6B, C) at SNI-8w. Notably, 7% of tdTomato^+^ neurons (2 out of 29) were SS type, and in contrast, a higher percentage, 18% (6/33), of tdTomato^-^ neurons were SS (Fig. 6C). Furthermore, we failed to observe any significant differences in RMP (Fig. 6D), Rheobase (Fig. 6E), AP firing rate (Fig. 6F, G), or AP firing rate in RS or LS neurons (Fig. 6H-K) comparing tdTomato^+^ vs tdTomato^-^ neurons in SNI-8w mice. These observations indicate that long-term chronic neuropathic pain causes marked changes in the intrinsic properties of different CeA neurons and perhaps also alters the local CeA microcircuit.

**Figure 6.**
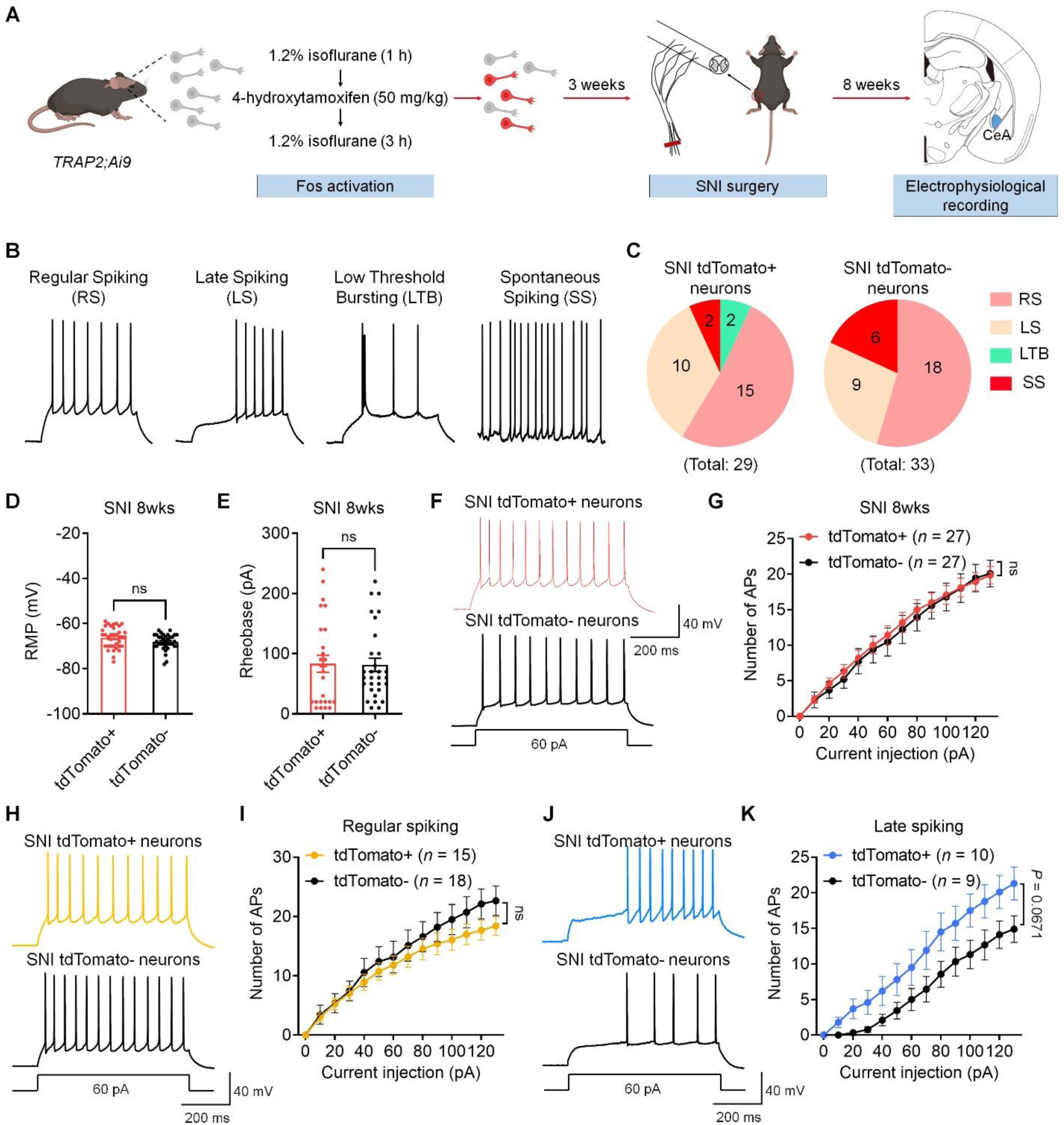
Comparison of firing patterns and excitability in CeA tdTomato^+^ neurons and tdTomato^-^ neurons in a chronic pain condition after SNI. **(A)** Schematic of experimental design for labeling CeA_GA_ neurons with tdTomato and electrophysiological recording in TRAP2; Ai9 SNI mice. **(B)** Representative voltage recordings of regular spiking (RS) neurons, late spiking (LS), low threshold bursting (LTB) and spontaneous spiking (SS) in within recorded tdTomato^+^ and tdTomato^-^ cell populations from TRAP2; Ai9 SNI mice. **(D)** RMP recorded in CeA tdTomato^+^ and tdTomato^-^ neurons from TRAP2; Ai9 SNI mice. **(E)** Rheobase evoked by step current injection were recorded in CeA tdTomato^+^ and tdTomato^-^ neurons from TRAP2; Ai9 SNI mice. **(F)** Representative AP traces evoked by 60 pA current in CeA tdTomato^+^ and tdTomato^-^ neurons from TRAP2; Ai9 SNI mice. **(G)** Number of APs evoked by step current injection in CeA tdTomato^+^ and tdTomato^-^ neurons from TRAP2; Ai9 SNI mice. **(H)** Representative AP traces evoked by 60 pA current in CeA tdTomato^+^ and tdTomato^-^ neurons from TRAP2; Ai9 SNI mice under regular spiking phenotype. **(I)** Number of APs evoked by step current injection in CeA tdTomato^+^ and tdTomato^-^ neurons from TRAP2; Ai9 SNI mice under regular spiking phenotype. **(J)** Representative AP traces evoked by 60 pA current in CeA tdTomato^+^ and tdTomato^-^ neurons from TRAP2; Ai9 SNI mice under late spiking phenotype. **(K)** Number of APs evoked by step current injection in CeA tdTomato^+^ and tdTomato^-^ neurons from TRAP2; Ai9 SNI mice under late spiking phenotype. All data are presented as the mean ± SEM and analyzed by Two-way ANOVA (G, I, K). ns: not significant.

To further investigate the SNI induced changes, we compared electrophysiological properties of TRAPed CeA_GA_ (tdTomato^+^) (Fig. 7A) to unlabeled neurons (tdTomato^-^, Fig. 7B) between naive and late-phase nerve injury (SNI-8w) conditions. In SNI-8w, tdTomato^+^ neurons exhibited no significant changes in RMP (Fig. 7C), Rheobase (Fig. 7D), and AP firing rate (Fig. 7E, F). Interestingly, tdTomato^-^ non-CeA_GA_ neurons, while had no significant changes in RMP (Fig. 7G), exhibited significantly reduced Rheobase (Fig. 7H), and elevated AP firing rate (Fig. 7I, J). Collectively, our findings suggest that long-term chronic neuropathic pain mainly increased the excitability of non-CeA_GA_ neurons.

**Figure 7.**
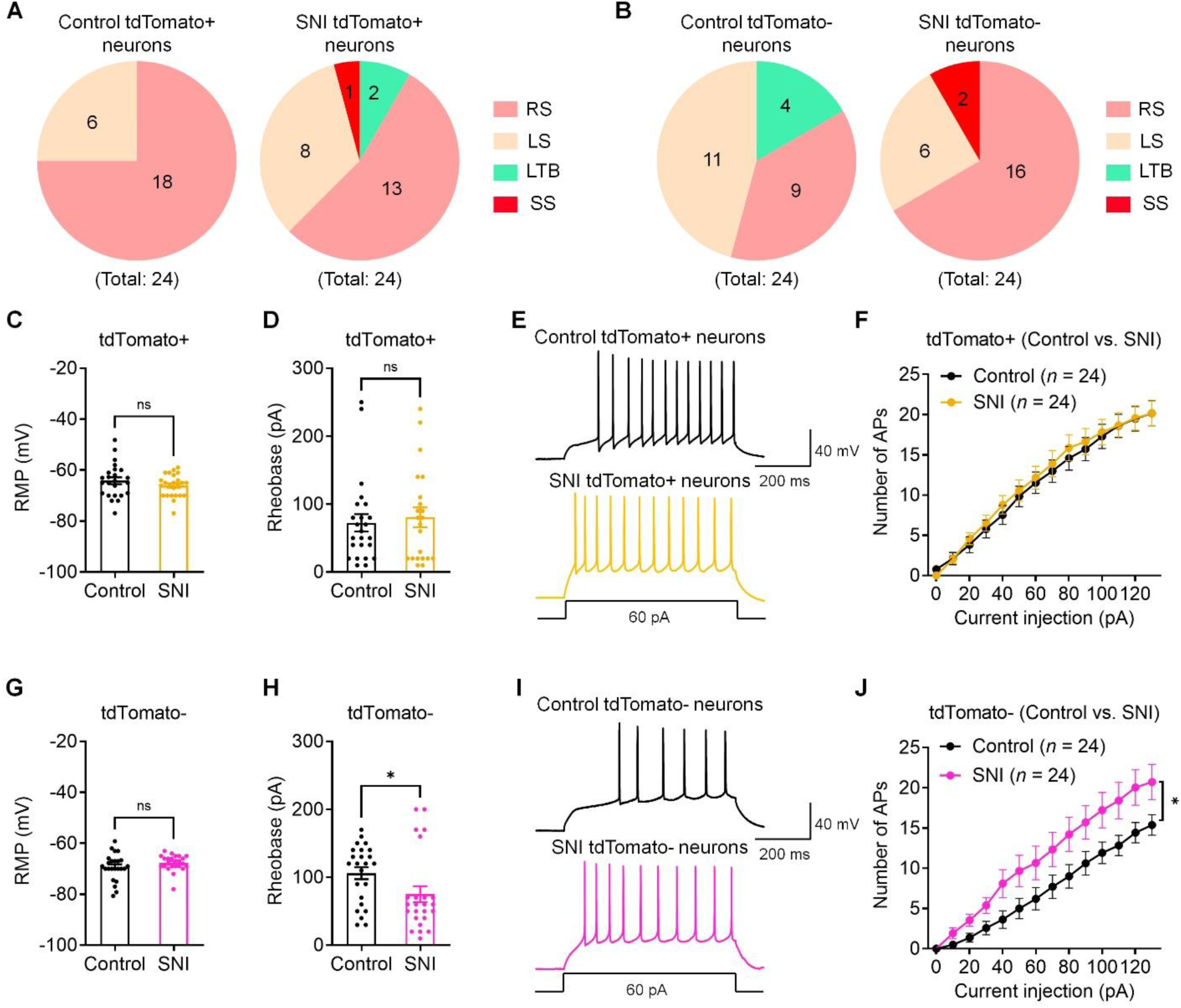
Chronic neuropathic pain (SNI-8weeks) enhances neuronal excitability of CeA tdTomato^-^ neurons but not tdTomato^+^ neurons. **(A)** Firing phenotype comparison of CeA_GA_ tdTomato^+^ neurons from control vs. SNI TRAP2; Ai9 mice. **(B)** Firing phenotype comparison of non-CeA_GA_ tdTomato^-^ neurons from control vs. SNI TRAP2; Ai9 mice. **(C)** RMP recorded in CeA_GA_ tdTomato^+^ neurons from control vs. SNI TRAP2; Ai9 mice. **(D)** Rheobase evoked by step current injection were recorded in CeA_GA_ tdTomato^+^ neurons from control vs. SNI TRAP2; Ai9 mice. **(E)** Representative AP traces evoked by 60 pA current in CeA_GA_ tdTomato^+^ neurons from control vs. SNI TRAP2; Ai9 mice. **(F)** Number of APs evoked by step current injection in CeA_GA_ tdTomato^+^ neurons from control vs. SNI TRAP2; Ai9 mice. **(G)** RMP recorded in tdTomato^-^ (non-CeA_GA_) neurons from control vs. SNI TRAP2; Ai9 mice. **(H)** Rheobase evoked by step current injection were recorded in tdTomato^-^ (non-CeA_GA_) neurons from control vs. SNI TRAP2; Ai9 mice. **(I)** Representative AP traces evoked by 60 pA current in tdTomato^-^ (non-CeA_GA_) neurons from control vs. SNI TRAP2; Ai9 mice**. (J)** Number of APs evoked by step current injection in CeA tdTomato^-^ (non-CeA_GA_) neurons from control vs. SNI TRAP2; Ai9 mice. All data are presented as the mean ± SEM and analyzed by unpaired two-tailed t-test (C, D, G, H) and Two-way ANOVA (F, J). \**P* < 0.05. ns: not significant.

### 3.6 CeA neurons exhibit distinct KCC2 expression patterns in both physiological and neuropathic pain states

It was known that KCC2 critically controls neuronal chloride homeostasis and maintains normal inhibitory synaptic transmission^20^. Down-regulation of KCC2 in spinal neurons after tissue and nerve injuries contributes to the transition from acute to chronic pain states^21–23^. These previous findings promoted us to examine KCC2 expression in CeA neurons under three different conditions (sham vs. early-phase SNI vs. late-phase SNI) in TRAP2; Ai9 mice (Fig. 8A-E). We found that KCC2 is preferentially expressed in tdTomato^-^ neurons (>90%) compared to tdTomato^+^ neurons (<10%, Fig. 8A, B). Nerve injury caused a significant KCC2 downregulation in the late-phase but not early phase of SNI (Fig. 8A, C). Further analysis showed that downregulation of KCC2 mainly occurred in non-CeA_GA_ tdTomato^-^ neurons during the transition from acute pain to chronic pain (Fig. 8A, E); and KCC2 expression in CeA_GA_ (tdTomato^+^) neurons remained low (Fig. 8A, D). These results suggest that the KCC2 downregulation in non-CeA_GA_ neurons might be responsible for the increased excitability of these neurons in chronic pain phase.

**Figure 8.**
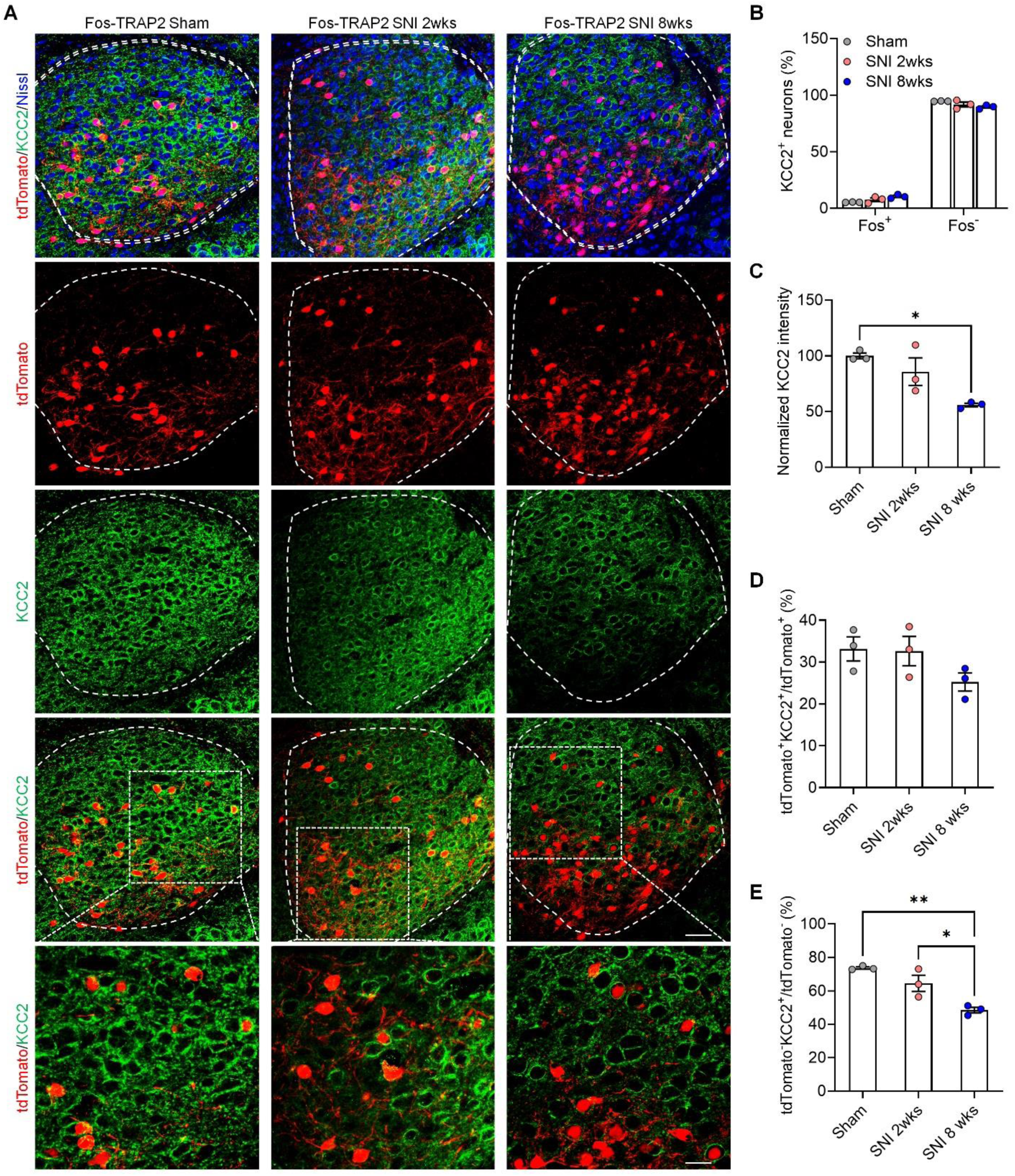
Distinct KCC2 expression alternations in CeA_GA_ tdTomato^+^ neurons and tdTomato^-^ neurons. (**A**) Representative KCC2 expression in CeA_GA_ tdTomato^+^ neurons and tdTomato^-^ neurons from sham, 2-week-SNI neurons over KCC2^+^ neurons in CeA regions from sham, 2-week-SNI and 8-week-SNI TRAP2; Ai9 mice. **(C)** Normalized KCC2 intensity in CeA regions from sham, 2-week-SNI and 8-week-SNI TRAP2; Ai9 mice. **(D)** Percentage of tdTomato^+^ KCC2^+^ neurons over tdTomato^+^ neurons in CeA regions from sham, 2-week-SNI and 8-week-SNI TRAP2; Ai9 mice. **(E)** Percentage of tdTomato^-^ KCC2^+^ neurons over tdTomato^-^ neurons in CeA regions from sham, 2-week-SNI and 8-week-SNI TRAP2; Ai9 mice. All data are presented as the mean ± SEM. \**P* < 0.05, \*\**P* < 0.01. One-way ANOVA, accompanied by Tukey’s post-hoc test for multiple comparisons.

## 4. Discussion

A previous study identified a specific subset of CeA neurons activated by general anesthesia that exhibited potent pain-suppressing effects^1^. Using an activity-dependent targeting method called CANE, this study revealed that optogenetic activation of these neurons reduced both mechanical and thermal pain sensitivity. In our current study, we validated the function of these neurons using an alternative method called Fos-TRAP2^6^. This method allowed us to successfully label the CeA_GA_ neurons activated during general anesthesia. Chemogenetic activation of the CeA_GA_ neurons resulted in significant pain-suppression effects in both naïve mice and mice with neuropathic pain (SNI model), especially in the early phase (SNI-2 weeks). In addition to validation, we also made several novel findings about CeA in different pain states. First, we characterized the electrophysiological properties of CeA_GA_ neurons and non-CeA_GA_ neurons in naïve mice, revealing that CeA_GA_ (tdTomato^+^) neurons exhibited higher excitability and distinct firing patterns compared to non-CeA_GA_ neurons. Second, we discovered that nerve injury induced a new firing pattern (spontaneous spiking) in subsets of both CeA_GA_ and non-CeA_GA_ neurons. Furthermore, in chronic pain states (SNI 8-weeks), non-CeA_GA_ neurons showed increased excitability. Third, we found that the analgesic effect of chemogenetic activation of CeA_GA_ neurons was potent in naïve mice and in mice with early-phase SNI (2 weeks) but was less effective in mice with late-phase SNI (8 weeks). Intriguingly, activation of CeA_GA_ neurons caused potent anti-anxiety effects during this late phase. Fourth, KCC2 was less expressed in CeA_GA_ neurons than in other CeA neurons, and its expression in non-CeA_GA_ neurons was downregulated in the late-phase SNI (8 weeks). Previous studies showed that different types of CeA neurons form local inhibitory connections among themselves^24,25^. Our data showed that activation of CeA_GA_ neurons have differential analgesic efficacies in acute versus late phase of chronic pain, and these differential effects were likely mediated by the altered intrinsic electrophysiological properties of CeA neurons and CeA microcircuit in part due to changes of KCC2 expression.

The previous study discovered CeA_GA_ as the forebrain central analgesic center, extending the pain modulation pathway beyond the midbrain periaqueductal gray-brainstem-spinal descending pathway^1,26–28^. CeA is a brain region consisting of different cell types, including PKCδ-, SST-, and Penk1-expressing neurons, among other types^29^. Using RNAcope method, we confirmed that the components of the CeA_GA_ neurons were 78.3% *Prkcd*^+^ neurons, 9.7% *Sst^+^* and 12% other neuronal cell types. In our study, we showed that chemogenetic activation of CeA_GA_ neurons attenuated pain in physiological and pathological conditions. Notably, study conducted by another group showed that the dual and opposing function of the CeA in pain modulation is encoded by opposing changes in the excitability of *Prkcd*^+^ and *Sst^+^* subpopulations of GABAergic neurons ^5^; and increased firing in *Prkcd*^+^ neurons enhanced pain-related responses while activation of *Sst^+^* neurons attenuated pain-related behaviors. It is possible that *Prkcd*^+^ neurons represent a much larger population of neurons than CeA_GA_ neurons, neurons within the first two weeks of nerve injury^5^, and the initial neuronal compensation for nerve injury in the early phase can be lost in the chronic pain.

The neuronal plasticity changes have critical roles in the modulation of CeA-dependent behaviors, including pain-related behaviors, fear conditioning and anxiety^5,24,30^. In our study, we characterized the electrophysiological properties of TRAPed CeA_GA_ and unlabeled neurons. These two subpopulations had distinct firing patterns: CeA_GA_ neurons showed greater neuronal excitability than non-CeA_GA_ neurons. This might contribute to CeA_GA_ neurons being activated by isoflurane in naïve mice. While both CeA_GA_ and other CeA neurons developed a new spontaneous spiking firing pattern in chronic pain condition, only non-CeA_GA_ neurons exhibited increased neuronal excitability. These results revealed that non-CeA_GA_ neurons had greater changes during the acute to chronic pain transition. Our observations that chemogenetic activation of CeA_GA_ neuron elicited potent pain-suppression in naïve mice and acute phase of nerve injury but weak pain suppression in chronic phase of neuropathic pain is consistent with a model in which analgesic CeA_GA_ and certain proalgesic non-CeA_GA_ neurons form reciprocal inhibitory microcircuits, and nerve injury elevated the excitability of non-CeA_GA_ neurons, thereby changing the balance of this inhibitory microcircuit and resulting in less effective CeA_GA_ mediated inhibition of proalgesic non-CeA_GA_ neurons, i.e. disinhibition, in chronic pain conditions. At present, we do not have a method to identify the critical proalgesic non-CeA_GA_ neurons to test this hypothesis.

Disinhibition in the spinal cord pain circuit is a general mechanism governing central sensitization and chronic pain and can be detected in the early phase of neuropathic pain^31–34^. This disinhibition in part is mediated by KCC2 downregulation in the spinal cord, since KCC2 plays a key role in maintaining the chloride ion gradient necessary for GABAergic inhibition in pain condition^8^. Interestingly, we found that chronic neuropathic injury also caused KCC2 downregulation in CeA, especially in non-CeA_GA_ neurons. The reduction in KCC2 might lead to a reversal of the chloride gradient^8^ and increased neuronal excitability, as well as impairment of the inhibition from analgesic CeA_GA_ neurons to proalgesic non-CeA_GA_ neurons in chronic pain conditions. Strategies such as increasing KCC2 expression or activation by pharmacological approaches ^9,35^, may restore the analgesic power of CeA_GA_ neurons for the management of chronic pain. Notably, even though chemogenetic activation of CeA_GA_ neurons was less effective in reducing hypersensitivity in late phase SNI, it still showed potent anti-anxiety effects, suggesting that different CeA_GA_ downstream target might be separately involved in modulating pain vs anxiety in chronic phase. In summary, our study validated the CeA_GA_ neurons as a central analgesic center and target the CeA_GA_ neurons and the underlying pathway could be a potential strategy to relieve acute and chronic pain.

## Acknowledgments

This study is supported by NIH grant R01DE29342 (F.W., J.P.M., R.R.J) and R01NS131812 (R.R.J)

## Author contributions

J.Z., F.W. and R.R.J. developed the project; J.Z., K.T, and A.M. conducted experiments and data analyses; J.Z. and R.R.J. wrote the manuscript; J.P.M., F.W. and other co-authors edited the manuscript; all authors approved the final manuscript submission.

## Declaration of interests

The authors have no completing financial interests in this study.

---

**Supplementary Figure 1.**
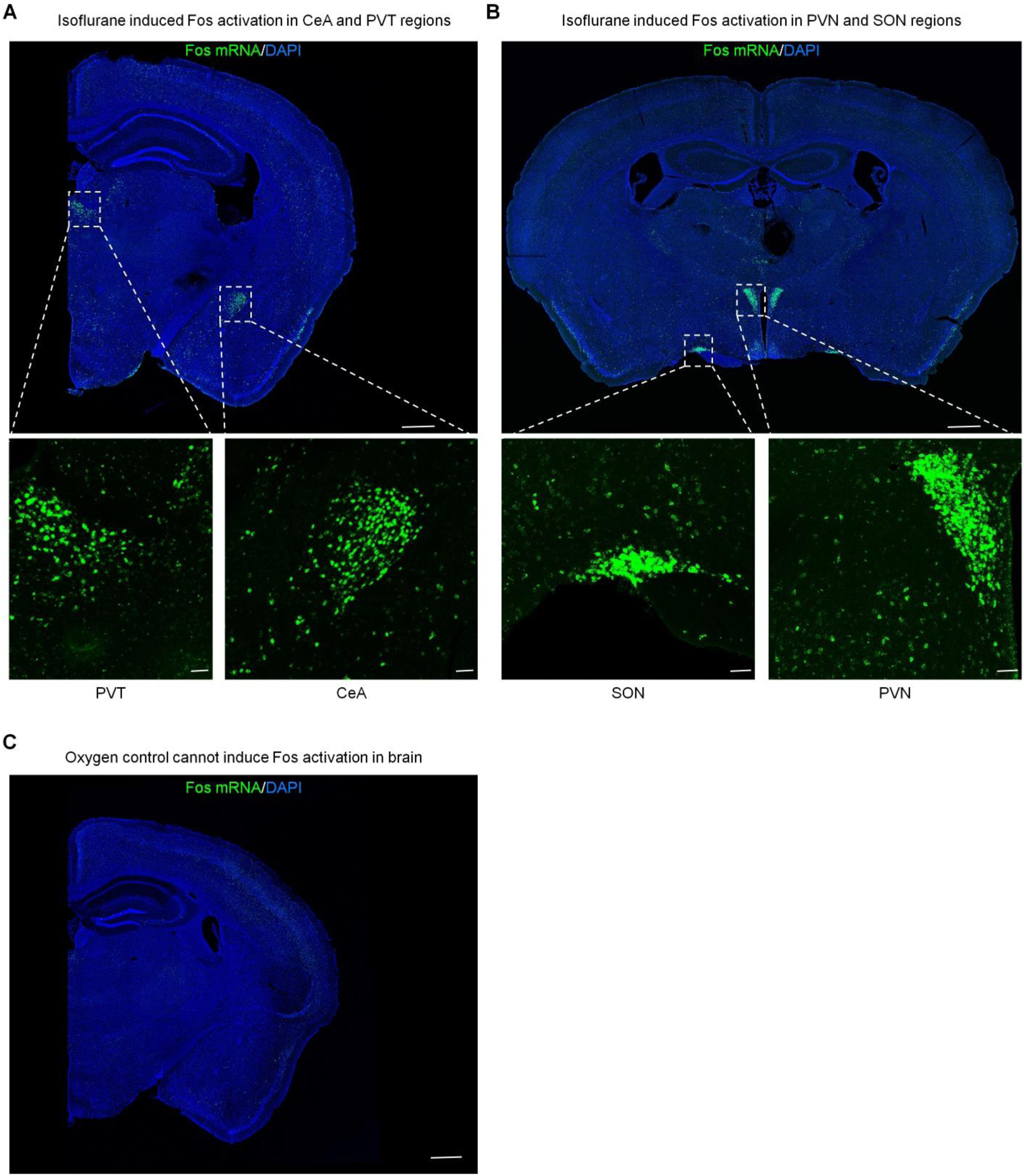
Isoflurane induces *Fos* expression in the CeA, PVT, PVN and SON of mouse brain. **(A)** Representative mouse brain image showing *Fos* induction in CeA and PVT regions after isoflurane exposure. **(B)** Representative mouse brain image showing *Fos* induction in PVN and SON regions after isoflurane exposure. **(C)** Negative control showing no *Fos* expression in CeA after oxygen exposure. Scale bars: 500 µm (low magnification in top panel) and 50 µm (high magnification).

## Notes

### Competing Interest Statement

The authors have declared no competing interest.

